# Multiphasic hepatitis B virus kinetic patterns in humanized chimeric mice can be explained via stochastic agent-based modeling of intracellular virion production cycles

**DOI:** 10.1101/2022.01.30.478385

**Authors:** Atesmachew Hailegiorgis, Yuji Ishida, Nicholson Collier, Michio Imamura, Zhenzhen Shi, Vladimir Reinharz, Masataka Tsuge, Danny Barash, Nobuhiko Hiraga, Hiroshi Yokomichi, Chise Tateno, Jonathan Ozik, Susan L. Uprichard, Kazuaki Chayama, Harel Dahari

## Abstract

Serum hepatitis B virus (HBV) kinetics in urokinase-type plasminogen activator/severe combined immunodeficient (uPA-SCID) mice reconstituted with humanized livers from inoculation to steady state is highly dynamic despite the absence of an adaptive immune response. We developed a stochastic agent-based model that includes virion production cycles in individual infected human hepatocytes. The model was calibrated using a genetic algorithm approach with the serum HBV kinetics observed in mice inoculated with 10^8^ HBV genome equivalents and fit the data well when the following viral production parameters were assumed: (1) An eclipse phase lasting 5-50 hours and (2) a post-eclipse phase production rate that is based on increasing production cycles initially starting with a long production cycle of 1 virion per 20 hours that gradually reaches 1 virion per hour after approximately 3-4 days before virion production increases dramatically to reach to a steady state production rate of 4 virions per hour per cell. The model was then validated by showing it could accurately simulate the viral kinetics observed with lower HBV inoculation doses (10^4^-10^7^ genome equivalents) in which similar, but delayed patterns were observed. Together, modeling suggests that it is the cyclic nature of the virus lifecycle combined with an initial slow but increasing rate of HBV production from each cell that plays a role in generating the observed multiphasic HBV kinetic patterns in humanized mice.

## Introduction

Despite the availability of an effective vaccine for the prevention of hepatitis B virus (HBV), HBV infection continues to impose an enormous burden with an estimated of 270 million chronically infected individuals and about 1 million deaths every year due to complications of HBV, including cirrhosis and liver cancer (1). Research to elucidate the molecular mechanisms that regulate the HBV life cycle and infection outcome (i.e., clearance vs. persistence) has been hampered by the lack of model systems that recapitulate HBV infection (2). Significant attempts have been made to develop small animal experimental models of HBV infection (3). The most successful small animal HBV infection model approach is based on liver-repopulation in immunodeficient mice with primary human hepatocytes as these are the natural target of HBV (4–8).

We previously assessed HBV infection kinetics from infection initiation to viral steady state in 42 chimeric urokinase-type plasminogen activator transgenic/severe combined immunodeficient (uPA-SCID) mice reconstituted with human hepatocytes (9). Serum HBV DNA was measured at varying intervals starting as early as 1 min post-inoculation and going out 63 days (9). Despite varying HBV doses (10^4^-10^8^ genome equivalents) and different batches of human hepatocytes, a consistent pattern of distinct phases was observed (9) (Fig.1). While mean-field mathematical models have been useful in understanding the dynamics of acute viral infection (10–18), these models were not designed to reproduce the multiphasic HBV infection kinetics we observed in these uPA-SCID mice (9).

Therefore, to elucidate the processes that result in the complex HBV kinetics observed from initiation to steady state in these mice, we have developed an agent-based modeling (ABM) approach that considers the cyclic nature of the viral life cycle within individual cells and the resulting distinct waves of viral release and spread. Specifically, the incorporation of viral production cycles within individual infected cells starting with an initially slow production (1 virion every 20 hours) that increases over time to reach steady state of 4 virions every hour recapitulates the multiphasic kinetic patterns. Using ABM to more accurate conceptualize HBV production as cycles rather than a continuous increase thus allows us to reproduce the observed HBV infection dynamics in vivo and provides a new modeling approach for simulating viral dynamics during acute infection.

## Results

### Agent-based modeling of HBV dynamics

While we showed (9) that the standard experimental approach of measuring HBV DNA levels in serial serum samples from infected mice allows for the calculation of average viral parameters and demonstrates complex multiphasic pattern of viral amplification from inoculation to steady state (Fig. 1), it masks the asynchronous infection of individual cells as the virus spreads (19). Because mean-field modeling approaches also do not take into account differences between individual cells during an asynchronous infection and cannot simulate the complex kinetics observed, we developed an ABM to investigate the dynamics underlying the observed multiphasic HBV kinetic pattern in humanized mice (Fig. 2). The ABM accounts for two types of agents: human hepatocytes (cells) and virus in blood. The cell agents, which are characterized by their infection stage are represented by a square lattice of 3 × 10^8^ cells i.e., the estimated number of human hepatocytes in the humanized mice (20). The cell agents can be in one of the following three discrete states: uninfected susceptible target (T), infected cell in eclipse phase (I_E_) (i.e., not yet releasing progeny virus), or productive infected cell secreting progeny virus (I_P_). The virus agent, V, represents the amount of HBV in blood and is characterized in the ABM as a single global agent. Due to the absence of an adaptive immune response in these mice, cell death is not considered in the model, hence the cell number was kept constant throughout the simulation period. The ABM execution is an iterative process where each iteration represents a “tick” or a discrete time step, where 1 step = 1 hour.

**Fig. 1.**
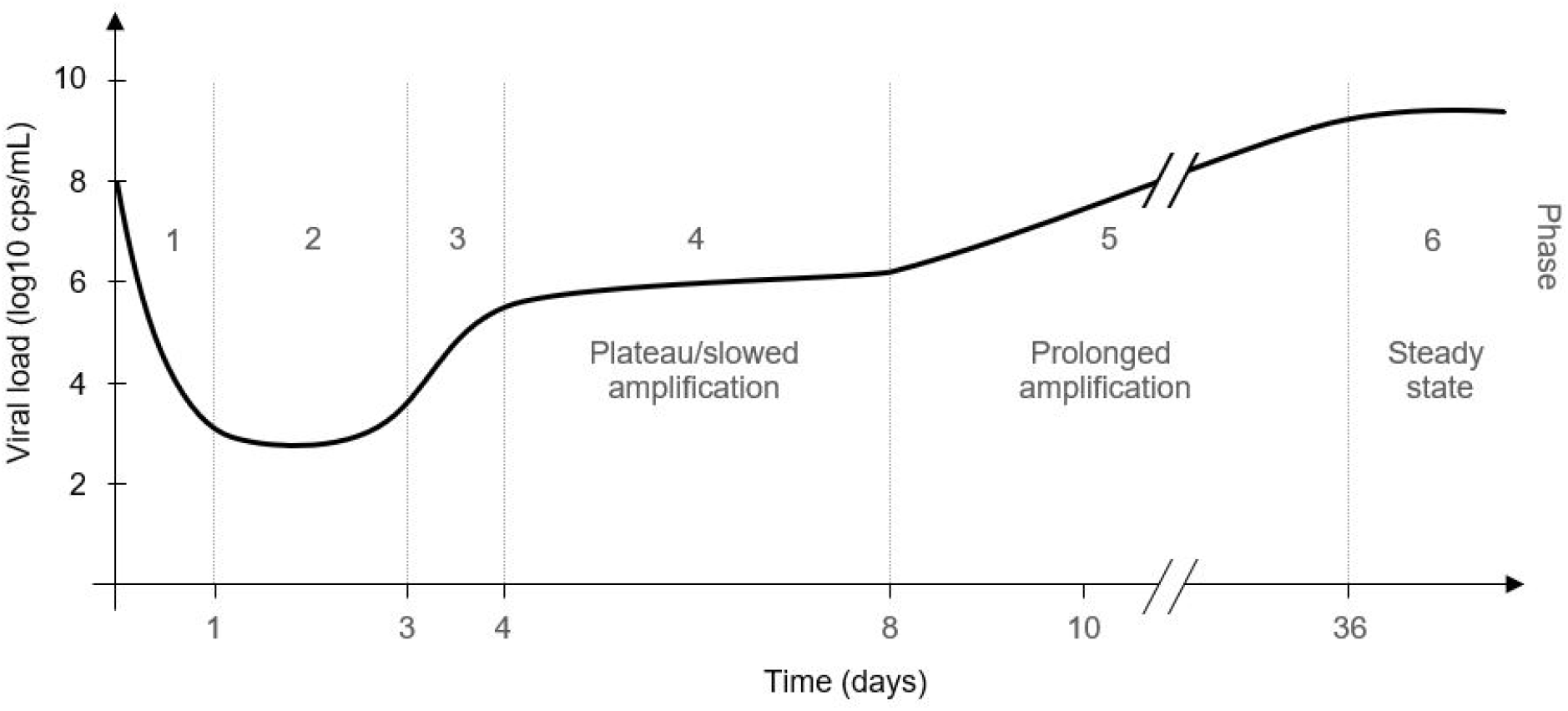
Overview of the main HBV kinetic patterns seen in humanized mice from inoculation to steady state. HBV kinetic phases from mice after inoculation with 10^8^ copies HBV DNA: Phase 1, rapid decline; Phase 2, lower viral plateau; Phase 3, rapid increase; Phase 4, extremely slow increase or plateau; Phase 5, prolonged amplification; Phase 6, steady state. Modified from (9); two initial clearance phases have been combined into one, now jointly termed Phase 1.

**Fig. 2.**
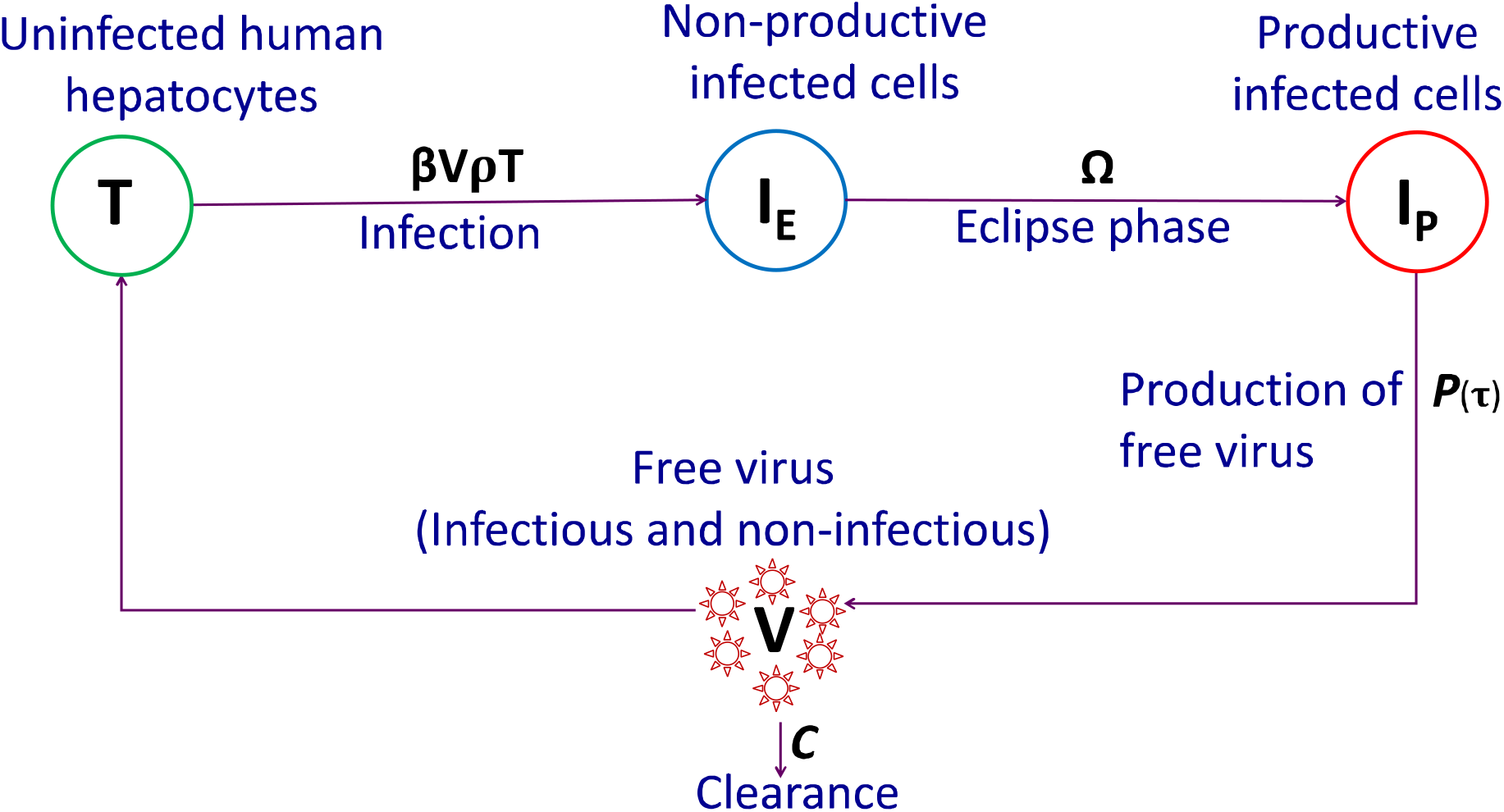
Schematic diagram of the ABM. The human hepatocytes can be only in one of the following three phases at a given time; T = uninfected cells which are termed as target or susceptible cells, I_E_ = HBV-Infected cells in eclipse phase (i.e., not yet releasing virions), I_P_ =productively HBV-infected cells (i.e., actively releasing virions). The free virus in blood, V, is composed of infectious and non-infections virions. The parameter ρ represents the fraction of virions that are infectious, β represents the infection rate constant, Ω represents eclipse phase duration, c, represents viral clearance from blood and P(τ) (Eq. 1) represents virion secretion from I_P_.

Viruses replicate exponentially and are obligate pathogens which use the host cell resources to replicate. Because HBV is a noncytolytic (21), chronic virus, viral production increases over time until reaching a resource restriction plateau. To model this on a per cell basis, we quantify both the amount of viral production by infected cells at a given time and the production cycle, i.e., the time interval over which the cell produces virus. To capture such dynamics, we formulated the following equations.

The amount of virion produced, P, at production cycle τ is determined as:

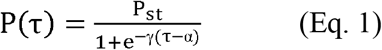

where, *P(τ)* is number of virions produced by infected cells a τ, *P_st_* is steady state virus production, α is number of cycles to reach to 50% of *P_st_*, γ is steepness of the production curve, and τ is the production cycle. Viral production, *P_st_*, is estimated at steady state when all target cells are infected, therefore *P_st_* = cV *_st_ /I _P_* where V_st_ represents viral load at steady state.

And the interval between cycles is determine by an exponential decay function with:

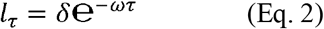

Where l_T_ is interval between production cycle, τ is the production cycle, δ is scaling factor indicating the initial production cycle length, and ω is decay constant. In the model, both P(τ) and l_T_ are roundup to the nearest integer. The combination of Eqs. 1 and 2 allows for each productively infected cell to slowly produce virions once I_E_ becomes I_P_ with increasing numbers of secreted virions within shorter time intervals until the cell reaches a steady state production.

### ABM reproduces multiphasic HBV kinetics after high dose inoculation

The model reproduces well the complex HBV DNA serum patterns observed in 4 mice inoculated with 10^8^ HBV genome equivalents and followed from inoculation until day 56 post-inoculation (p.i.)(Fig. 3). The estimated model parameters based on best fit for the 4 mice (M1-M4) are shown in Table 1. A graph showing the kinetics of the changing cell states (Fig. 4A) and representative pictures of the cell populations at distinct time points (Fig. 4B) during the simulation of infection for mouse 1 (M1) reveals that the progression of infection at the level of productively infected cell number (Fig. 4A, red curve) is not complex.

**Table 1.**
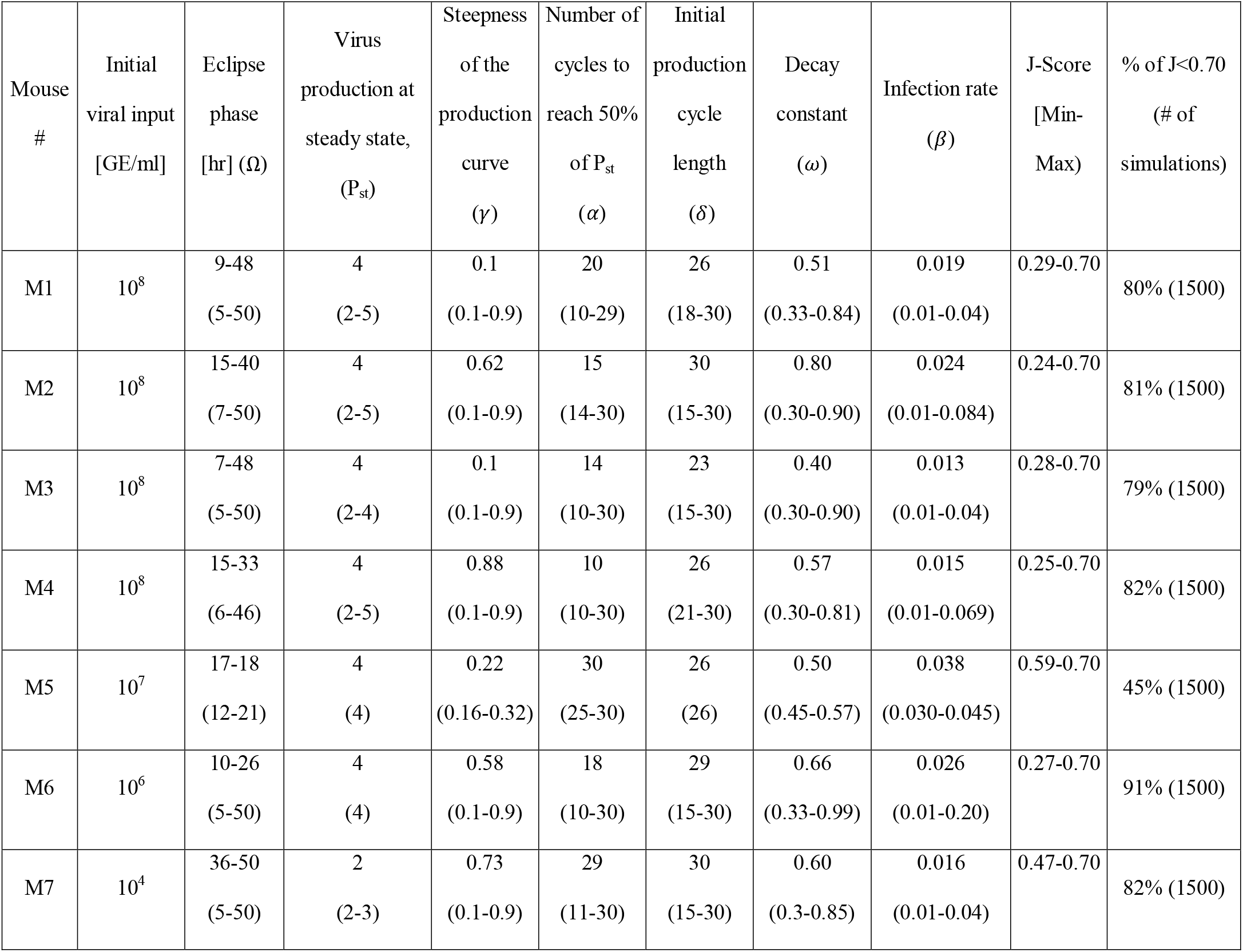
Best estimated ABM parameters and their estimated space (J<0.7): Parameters are defined in Eqs. 1 and 2; GE, genome equivalent; J-score, represents the objective function score of genetic algorithm (GA) fits where min score is the best fit curves shown in Fig. 3.

**Fig 3.**
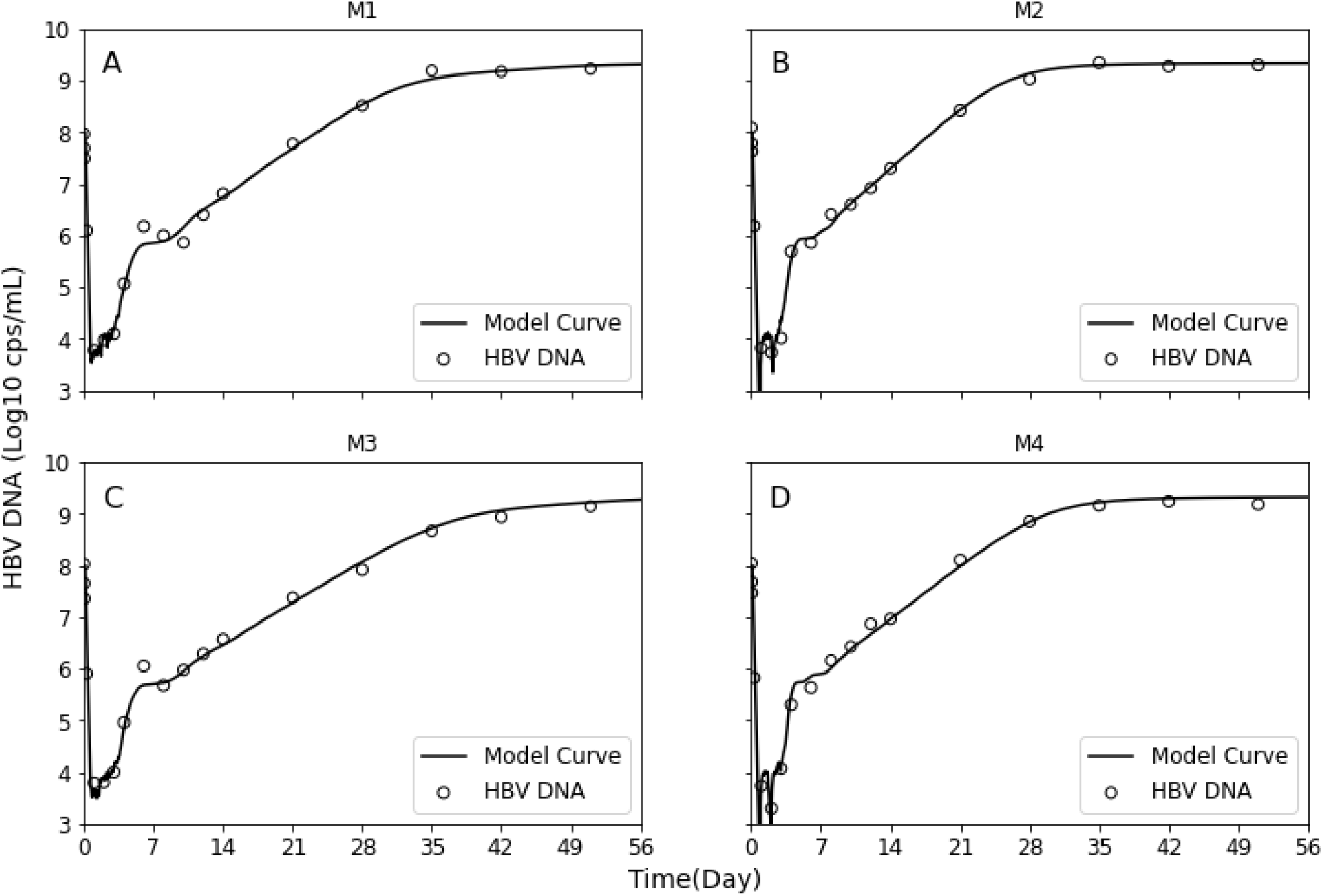
Model best fit (solid curves) with measured HBV DNA kinetics in blood (circles) in four mice M1 **(A)**, M2 **(B)**, M3 **(C)** and M4 **(D)** inoculated with 10^8^ HBV genome equivalents. Estimated parameters are shown in Table 1.

**Fig. 4.**
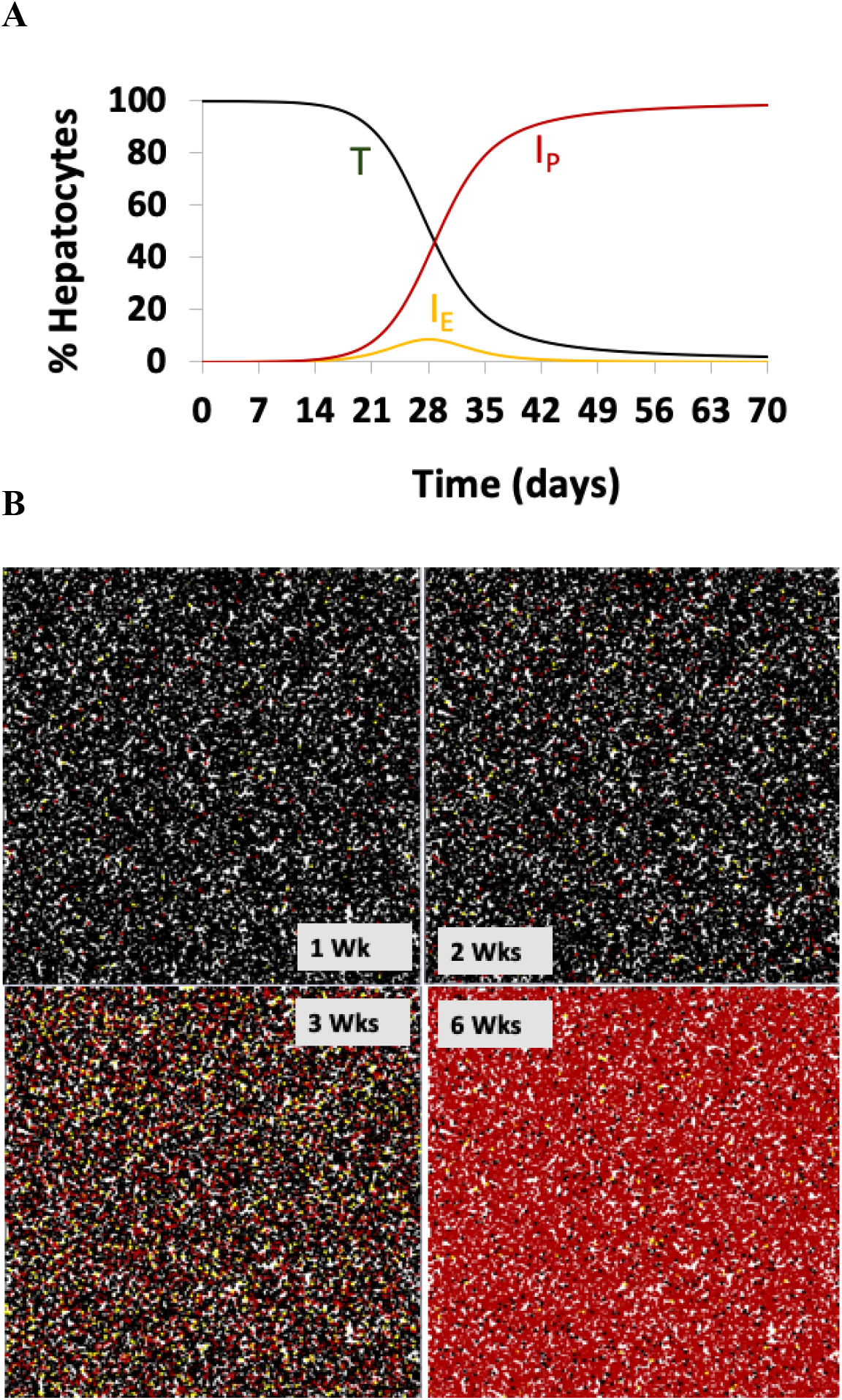
Human hepatocytes infection kinetics. (**A**) kinetics of cell infection status. (**B**) kinetics at 1, 2, 3, and 6 weeks post inoculation in a lattice. Uninfected cells (T, black line/cells), Infected non-productive cells (I_E_, yellow line/cells), and productively-infected (I_P_, red line/cells) cells. The ABM results represent the best fit of mouse M1 (Fig. 3A, Table 1).

### Virus parameter estimates of individual cells that allow for simulation of the observed complex serum viral patterns

The model provides insights into early virus-host dynamics of individual cells from infection to steady state that unmask the nature of the observed complex serum viral kinetic patterns. To fit the data, the model predicts a variable eclipse phase ranging from 5-50 hours in the 4 mice inoculated with 10^8^ HBV genome equivalents (Table 1). The simulated virion production for M1 with two eclipse phase durations of 9 hr (Fig. 5A, shaded box) and 48 hr (Fig. 5C, shaded box) illustrates the impact this has on the kinetics of virion production. Post-eclipse phase, the model predicts viral release from productively infected cells starts slowly with a long production cycle of 1 virion per 20 hours that gradually reaches 1 virion per hour (Fig. 5 B and D) after ~3-4 days before virion production increases to reach to a steady state production rate of 4 virions per hour per cell (Fig. 5 A and C).

**Fig 5.**
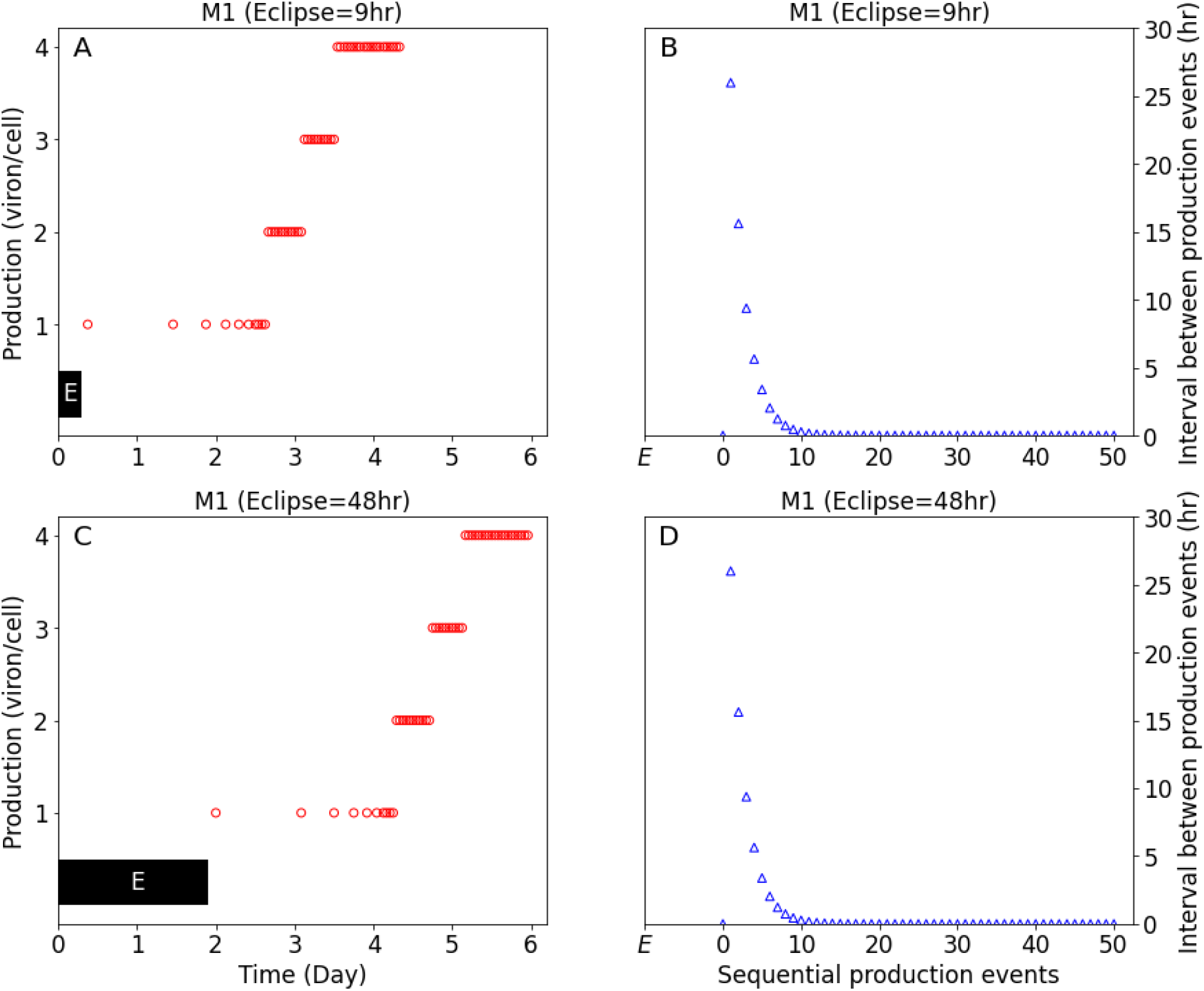
HBV virion production (circles) and viral production cycles (triangles) in representative mouse (M1) inoculated with 10^8^ HBV genome equivalents shown in Fig. 3A. (**A** and **B**) represent infected cells with minimum eclipse phase of 9 hr (black-shaded “E”). (**C** and **D**) infected cells with maximum eclipse phase of 48hr.

### Dissecting the nature of each serum HBV DNA kinetic phase

Focusing on the details of the model simulation for representative M1, the estimated serum HBV DNA is shown on a time scale that allows visualization each serum HBV DNA kinetic phase (Fig. 6 A and B). The number of non-productive (I_E_) and productive (I_P_) infected cells along with the average virion production per productive cells are plotted on the same scale for direct comparison (Fig. 6 C and D).

**Fig. 6.**
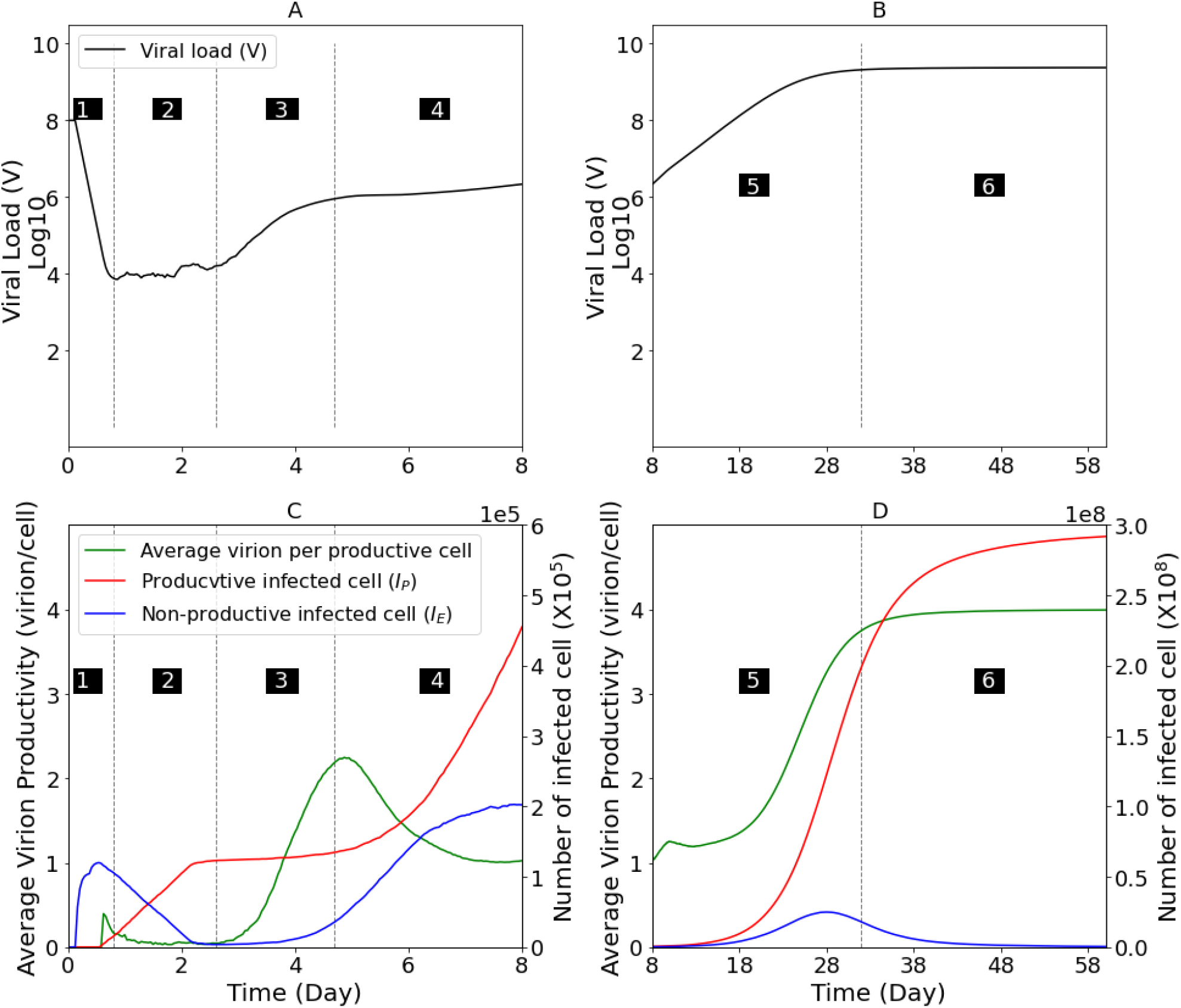
Model parameter estimates for representative mouse M1. (**A and B**) Simulated serum viral load. (**C and D**) The number of total cells in eclipse phase (blue line), productively infected cells (red line) and the average virion production (or secretion) per productively infected cells (green line) per time post inoculation. Graphs are divided according to the kinetic phases of HBV serum DNA amplification observed experimentally in Figure 1. Dashed vertical lines and black-shaded numbers indicating kinetic Phases 1 – 4 in (**A**) and (**C**) and Phases 5 and in (**B**) and (**D**).

During the first 6 hours p.i. (Fig. 6A, Phase 1), as the model recapitulates the rapid serum HBV DNA (V) clearance from the blood (t_1/2_=1 hr), it predicts that about 1×10^5^ cells were infected, i.e., the first wave of infection. This consists of an initial peak of cells of eclipse phase cells (I_E)_ (Fig. 6C, blue line).

As the I_E_ gradually transition into I_P_ (Fig. 6C, red line) the resulting low initial virion production (Fig. 6C, green line), balances continual viral clearance resulting in the lower serum viral plateau (Fig. 6A, Phase 2).

The first rapid increase in HBV serum levels (Fig. 6A, Phase 3) occurred once the virus production rate increases in the initially infected I_P_ cells (Fig. 6C, green line). During Phase 3 the number of I_P_ cells remained constant (Fig. 6C, red line) while the second wave of newly infected I_E_ cells start to emerge (Fig. 6C, blue line).

The observed intermediary serum HBV DNA steady state (or plateau) is recapitulated by the model (Fig. 6A, Phase 4) as the majority of I_P_ cells are at steady state levels of viral production and the new increasing numbers of I_P_ cells are in the early low level virus production phase of infection (Fig. 6C, increasing red line). Likewise, the increasing number of I_E_ cells do not contribute to virion production (Fig. 6C, increasing blue line) resulting in an overall decrease in the per cell virion production rate (Fig. 6C, decreasing green line).

As the infection becomes less synchronized due to the stochasticity that exists in the timing of individual infection events and target cells become limiting, the distinct cycles of infection become less discernable and all subsequent amplification appears as a single viral exponential expansion from day ~8 until ~30 days p.i. (Fig. 6B, Phase 5) during which the final target cells are infected and become productive (Fig. 6D, blue and red lines, respectively) and subsequently progress towards maximal average viral production (Fig. 6D, green line).

Once all of the target cells are productively infected (Fig. 6B, red line) and achieve maximal average viral production (Fig. 6D, green line) serum HBV levels attained steady state (Fig. 6B, Phase 6).

### ABM reproduces multiphasic HBV kinetics after low dose infection

We previously showed that lower inoculation of 10^7^, 10^6^ and 10^4^ HBV genome equivalents in humanized uPA-SCID mice led to a delayed, but similar complex HBV kinetic pattern (9). Therefore, we validated the model against this kinetic data by changing only the inoculation dose in the model simulation. Importantly, the model reproduced well the kinetic patterns within the same parameter space estimated for mice that were inoculated with 10^8^ HBV genome equivalents (Fig. 7).

**Fig. 7.**
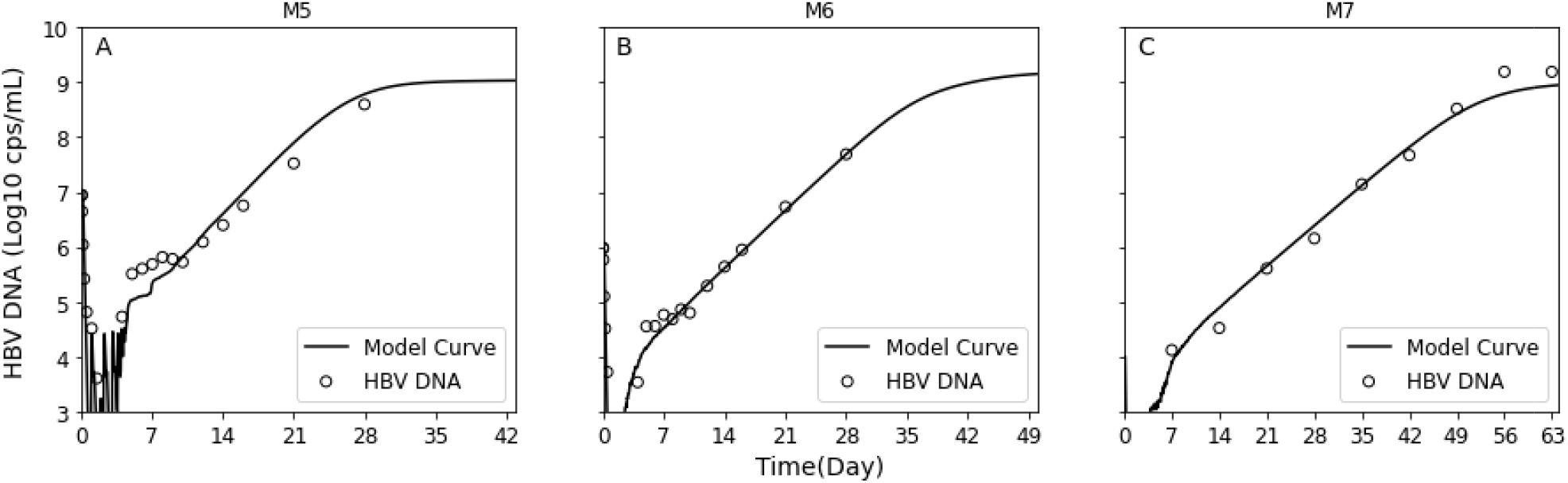
Model validation. Model best fit curves (solid lines) with measured HBV DNA kinetics (circles) in mice M5, M6 and M7 inoculated with and 10^7^ **(A)**, 10^6^ **(B)**, and 10^4^ **(C)** HBV genome equivalents, respectively. Estimated model parameters are shown in Table 1.

### Short virion production cycle and eclipse phase lengths diminish multiphasic kinetic pattern

Having validated the ability of the model to accurately recapitulate the kinetics of HBV infection, we proceed to investigate how the two key features of the model, namely cellular eclipse phase and/or production cycle, might affect acute viral infection kinetics in general. To test the effect of shortening and/or lengthening the cellular eclipse phase and/or production cycle, in silico simulations were performed using HBV viral parameters estimated for mouse 1 (M1) as a reference for 14 days (Fig. 8) post inoculation.

**Fig 8.**
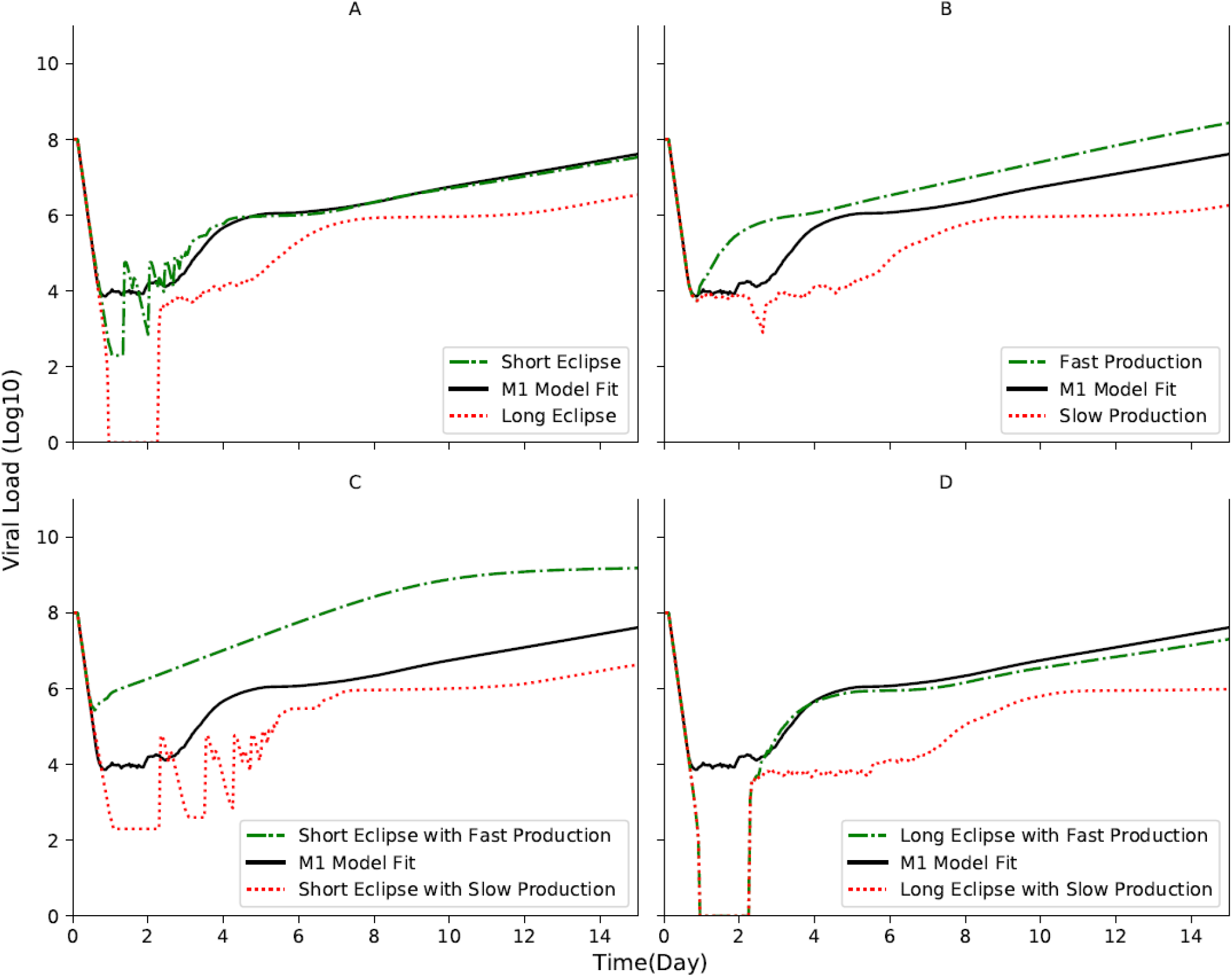
Varying the eclipse phase length (Ω) and initial production cycle length (O). Model simulations were run from time of inoculation until day 14 post inoculation (p.i.). Parameters equal to that estimated for mouse 1 (M1) were used except for the indicated changes. **(A)** The parameter range of the eclipse phase was shortened to Ω = [0,5] hr (dashed green line) or extended to Ω =[36-72] hr (dotted red line). **(B)** The parameter range for the production cycle was reduced to δ =1 hr (i.e., faster production, dashed green line) or increased to δ = 36 hr (i.e., slower production, dotted red line). **(C)** The short eclipse parameter range of Ω = [0,5] hr was combined with the fast (dashed green line) or slow (dotted red line) production parameter ranges used in (B). **(D)** same as **(C)** assuming extended eclipse phase to Ω =[36-72] hr. The model simulations for M1, Ω = [9,48] hr and to δ = 26 hr, is shown for comparison using solid black lines.

Lengthening the eclipse phase resulted in an enhanced initial lower plateau and followed by a delayed but multiphasic (Phases 2-6) viral amplification (Fig. 8A, red dotted line). In contrast, a short eclipse phase allowed for visualization of more extreme cycling during Phase 2 of the infection without affecting subsequent viral phases (Fig. 8A, green dashed line). Increasing the virion production resulted in a much shorter lower viral plateau phase (Phase 2) and earlier amplification with a less pronounced Phase 2 (Fig. 8B, green dashed line). Slower production in contrast lengthened the initial lower plateau phase, Phase 2, and delayed subsequent viral phases (Fig. 8B, red dotted line). Interestingly, combining a shorter cellular eclipse with a faster production cycle produces an almost single amplification phase analogous to that observed for many acute viral infections (Fig. 8C, green dashed line) perhaps suggesting why such multiphasic viral infection kinetic patterns have not been universally observed. In contrast, combining a shorter cellular eclipse with a slower production cycle produces again the short eclipse extreme cycling this time during a longer Phase 2 followed by delay of all subsequent viral phases (Fig. 8C, red dotted line). Combining a longer cellular eclipse with a faster production cycle resulted in an enhanced lower plateau (Phase 2) with little effect on subsequent viral phases (Fig. 8C, green dashed line). However, when a longer cellular eclipse is coupled with a slower production cycle, the enhanced initial lower plateau is followed by delayed but multiphasic (Phases 2-6) viral amplification (Fig. 8D, red dotted line).

## Discussion

Standard population-based measurements of viral infection time courses typically show an initial eclipse phase followed by exponential increase ending either in cell lysis or steady state viral levels (10–18). However, we recently reported a surprising complex HBV infection kinetics in chimeric uPA-SCID mice with humanized livers (9). To investigate the dynamics underlying the unexpectedly complex HBV infection kinetics from inoculation to steady state in humanized uPA-SCID mice, we developed an ABM approach to explain the serum population kinetics from the perspective of individual cell infections. This allowed for the simulation of HBV production from individual cells as cycles rather than a continuous increase which resulted in the model accurately reproducing the multiphasic kinetic pattern observed.

The majority of mathematical models of acute viral infection are based on the assumption of well-mixed virus and cell populations (10–17). In the absence of the development of immune response in which infected cells will not lost or die in a faster rate compared to uninfected cells, these models predict a roughly monophasic viral increase that will reach a high viral load steady state once all target cells are infected. Some models also accounted for an eclipse phase (e.g. (22, 23)), in which newly infected cells remain in a latent phase before becoming virion producing infected cells, but even with such this addition these models cannot predict multiphasic viral kinetics in the absence of immune response. For example, in the chimpanzee model of acute hepatitis C virus (HCV) infection, viral levels increased in a biphasic manner with a transient viral decline in between (1 week p.i.) concomitantly with the induction of type I interferon (13). However, HBV is often referred to as a stealth virus because it does not induce significant innate immune signaling (21), making immune signaling a less likely explanation for the shift from rapid expansion (Fig. 1, Phase 3) to a slower interim plateau (Fig. 1, Phase 4). While intrahepatic interferon induction cannot be ruled out without further experiments, we show here that this shift as well as all the phases observed can be accurately recapitulated by describing viral dynamics at the individual cell level combining the viral eclipse phase, increasing rate of virus secretion based on increasing production cycles, and the resulting waves of new infection (Fig. 6).

Viral kinetics early post-infection often exhibits a viral decline phase followed by lower plateau or undetectable viral level (a.k.a. eclipse phase) before exponential amplification. This is assumed to reflect the time period in which clearance of input virus in the blood is balanced by de novo viral production. We showed that mice inoculated with high (10^8^) virus inoculation HBV DNA, had a low viral plateau which lasted approximately 1-3 days p.i (Fig. 1 Phase 2). Fitting the ABM with measured serum HBV DNA we estimated a range of 5 and 50 hours in which newly infected cells (I_E_) cell could rest in a latent phase before becoming virion producing infected cells (I_P_). While this initially seemed like an unexpectedly broad parameter range, recent single cell analysis of viral infections has revealed analogously large ranges of variability in the progression of replication (24, 25).

We previously reported (9) that HBV DNA inoculum size had no effect on initial HBV clearance (Fig. 1, Phase 1), the viral doubling time during the HBV expansion phases (Fig. 1, Phases 3 and 5), the length of the interim plateau (Fig. 1, Phase 4), or viral steady-state levels (Fig. 1, Phase 6), but rather resulted in a lower viral plateau (Phase 2) and a delay in detection of initial virus expansion (Fig. 1, Phase 3), which subsequently delayed all other kinetic phases. Importantly, the model confirms this as simulating the infection kinetics in mice after HBV inoculation of 10^7^, 10^6^ or 10^4^ genome equivalents, requires no significant change in any of the estimated ABM parameters compared to the mice inoculated with 10^8^ genome equivalents except for the inoculation dose itself (Table 1), further supporting our ABM modeling approach.

Finding that the unusual multiphasic kinetics observed can be reproduced by a model that is based on the inherent cyclic nature of a viral life cycle, raised the question why such complex kinetics was observed for HBV and not more broadly for other viral infections. While it is possible that such a multiphasic pattern could simply be missed in the absence of frequent sampling, this complex pattern was also eliminated by increasing viral production and reducing the eclipse phase in the model simulations (Fig. 8C). Notably, these parameter changes are consistent with the faster lifecycles associates with many acute viruses and may explain why multiphasic viral kinetics has not been routinely observed for other viruses..

The current ABM does not account for intracellular HBV-host dynamics that explain the estimated timing of the viral production cycles but provides a prediction that can now be investigated. Our previous kinetic study (9), indicated that intrahepatic total HBV DNA, cccDNA, and RNA correlate with serum HBV DNA during infection in these mice. Thus, a future detailed intracellular kinetic analysis may shed light whether intrahepatic HBV RNA and/or DNA levels exhibit the same multiphasic amplification pattern and how cccDNA recycling impacts the level of HBV secretion early in infection.

Meanwhile, we show in the current study that the incorporation of viral production cycles into a stochastic ABM recapitulates the multiphasic serum HBV kinetic patterns observed in uPA-SCID mice reconstituted with humanized livers from inoculation to steady state (9). Importantly, such complex HBV kinetic infection can be seen also in immunocompetent chimpanzees (11, 26) indicating that this complex picture is not unique to humanized uPA-SCID mice. Thus, using agent-based modeling to more accurate conceptualize virus production as cycles rather than a continuous increase allows us to reproduce the observed HBV infection dynamics in vivo and provides a new computational approach for simulating viral dynamics during acute infection.

## Methods

### Mice and HBV

Humanized liver chimeric mice were produced by splenic injection of cryopreserved human hepatocytes into uPA/SCID mice as previously described (8, 9). Human hepatocyte repopulation rates were estimated by blood human albumin levels and chimeric mice showing higher repopulation rates (>70%) were injected with serum containing 10^4^, 10^6^, 10^7^ and 10^8^ copies HBV DNA (genotype C, Accession No. AB246345, originally provided by Dr. Sugiyama (27) via the tail vein. Preparation of the inoculum, DNA extraction and quantification of HBV DNA were performed as previously reported (9, 28). As previously reported (9), all animal protocols from which the data in this manuscript were derived were performed in accord with the Guide for the Care and Use of Laboratory Animals and approved by the Animal Welfare Committee of Phoenix Bio Co., Ltd.

### Parameter estimations

We previously showed that during the first 6 h p.i., HBV was cleared from blood with a t_1/2_ of ~1 h independent of inoculum size which ranged from 10^4^ to 10^8^ cp/ml (9). Thus, we assumed a fixed clearance rate of virus from blood as c=0.67 ±0.09 h^−1^. The fraction of virus in the blood (V) that was infectious was arbitrary set as ρ=0.5. Based on the experimental data, HBV serum levels at steady state, V_St_ = 9.3±0.3. Viral production, P_St_, is estimated at steady state where all target cells are infected (I_P_ =3×10^8^ cells), which is equivalent to 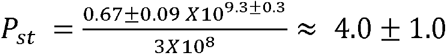 virion/cell. The remaining parameters were estimated by calibrating the ABM with the experimental data (Table 1).

### Model calibration

Model parameter fitting was done using a Genetic Algorithm (GA) (29, 30) with the EMEWS framework (31) on the Midway2 high-performance computing (HPC) cluster at the University of Chicago. Midway2 has 400 nodes, each with 28 cores and 64GB of memory. Some additional development was done on the Bebop HPC cluster, managed by the Laboratory Computing Resource Center at Argonne National Laboratory. Bebop has 1024 nodes comprised of 672 Intel Broadwell processors with 36 cores per node and 128 GB of RAM and 372 Intel Knights Landing processors with 64 cores per node and 96 GB of RAM. The GA was implemented using the DEAP (32) evolutionary computation Python framework (specifically(33): Chapter 7) and integrated into an EMEWS HPC workflow using EMEWS queues for Python (EQ/Py) (31). The use of HPC resources enable the concurrent evaluation of large numbers of design points (102), reducing the time to solution. During each iteration of the GA, the best points from the currently evaluated population are selected using a tournament selection method to create a new population. Each of these points is then mated with another according to a crossover probability and, finally, each of the resulting points is mutated according to a mutation probability. At each GA algorithm iteration, the new population is evaluated in parallel and the evaluation results are gathered. The GA population size was set to 102, the mutation probability to 0.2, the crossover probability to 0.5, and the number of iterations to 25. On Midway2 the runtime for a typical run was 7.3 hours using full concurrency on 28 nodes (with 28 cores per node), or about 5700 core hours.

## Acknowledgments

This work was supported by National Institutes of Allergy and Infectious Diseases Grants R01AI144112 and R01AI146917 and Japan Agency for Medical Research and Development (AMED) under grant 19fk0210020h0003, the Fund for the Promotion of Joint International Research (Fostering Joint International Research) from Japan Society for the Promotion of Science (Grant number: 17KK0194). This work was completed in part with resources provided by the Research Computing Center at the University of Chicago (the Midway2 cluster), and the Laboratory Computing Resource Center at Argonne National Laboratory (the Bebop cluster).

## Notes

### Competing Interest Statement

The authors have declared no competing interest.

